# Fe(III) heme sets an activation threshold for processing distinct groups of pri-miRNAs in mammalian cells

**DOI:** 10.1101/2020.02.18.955294

**Authors:** Sara H. Weitz, Jen Quick-Cleveland, Jose P. Jacob, Ian Barr, Rachel Senturia, Kikuye Koyano, Xinshu Xiao, Shimon Weiss, Feng Guo

## Abstract

The essential biological cofactor heme is synthesized in cells in the Fe(II) form. Oxidized Fe(III) heme is specifically required for processing primary transcripts of microRNAs (pri-miRNAs) by the RNA-binding protein DGCR8, a core component of the Microprocessor complex. It is unknown how readily available Fe(III) heme is in the largely reducing environment in human cells and how changes in cellular Fe(III) heme availability alter microRNA (miRNA) expression. Here we address the first question by characterizing DGCR8 mutants with various degrees of deficiency in heme-binding. We observed a strikingly simple correlation between Fe(III) heme affinity *in vitro* and the Microprocessor activity in HeLa cells, with the heme affinity threshold for activation estimated to be between 0.6-5 pM under typical cell culture conditions. The threshold is strongly influenced by cellular heme synthesis and uptake. We suggest that the threshold reflects a labile Fe(III) heme pool in cells. Based on our understanding of DGCR8 mutants, we reanalyzed recently reported miRNA sequencing data and conclude that heme is generally required for processing canonical pri-miRNAs, that heme modulates the specificity of Microprocessor, and that cellular heme level and differential DGCR8 heme occupancy alter the expression of distinct groups of miRNAs in a hierarchical fashion. Overall, our study provides the first glimpse of a labile Fe(III) heme pool important for a fundamental physiological function and reveal principles governing how Fe(III) heme modulates miRNA maturation at a genomic scale. We also discuss potential states and biological significance of the labile Fe(III) heme pool.

## INTRODUCTION

Heme is a complex of iron and protoporphyrin that is essential for life. Most iron in the human body is present as part of heme. Heme participates in numerous biological pathways, by serving as a prosthetic group for heme proteins, or as a signaling molecule to regulate cellular processes (Ponka 1999; Tracz et al. 2007; Zhang 2011). For cells to function normally, a fine balance is maintained between heme synthesis and degradation through feedback and feedforward mechanisms (Ryter and Tyrrell 2000; Ye and Zhang 2004; Khan and Quigley 2011). Over-abundance of heme causes severe cell and tissue damages, and increased heme synthesis and uptake are involved in cancer (Hooda et al. 2014). Conversely, heme deficiency is central to anemia and porphyria. For normal physiological functions, heme is also transported between organs, cells, and subcellular compartments (Reddi and Hamza 2016).

Most heme molecules in cells and tissues are already bound to their host proteins. However, it is the pool of labile heme, heme molecules that have not reached their destinations, that determines whether newly synthesized heme proteins are loaded, and whether a signaling pathway is turned on or off. Labile heme in cells is transient and is dynamically regulated, leading to technical challenges in detection and functional studies (Sassa 2004). Based on effects of heme in the expression of heme synthesis and degradation enzymes and substrate affinities of the heme degradation enzymes, heme oxygenases, it was proposed that the concentration of labile heme may be around 100 nM (Granick et al. 1975). Recently, three labile-heme detection methods have been developed (Song et al. 2015; Hanna et al. 2016; Yuan et al. 2016). They showed that labile heme in cells is compartmentalized. The estimated concentrations are in the nanomolar range with values depending on cell types and methods used. Two of these heme sensors favor binding Fe(II) heme (Song et al. 2015; Hanna et al. 2016) and the redox specificity of the third one is unclear (Yuan et al. 2016). It is currently unknown how much labile heme is in the Fe(III) form. It has been suggested that labile heme molecules are bound to heme transporter or chaperone proteins in cells (Severance and Hamza 2009).

Heme activates miRNA maturation by binding to DGCR8 (*D*i*G*eorge *c*ritical *r*egion gene *8*) (Faller et al. 2007; Barr et al. 2012; Weitz et al. 2014). DGCR8 and the ribonuclease III Drosha form the Microprocessor complex, which recognizes and cleaves pri-miRNAs to produce processing intermediates called precursor miRNAs (pre-miRNAs) in the nucleus as the first step of the miRNA maturation pathway (Denli et al. 2004; Gregory et al. 2004; Han et al. 2004; Landthaler et al. 2004). DGCR8 contains a dimeric RNA-binding heme domain (Rhed, residues 276-498 in humans) that binds to the apical end of pri-miRNA hairpins and works with the two double-stranded RNA-binding domains (dsRBDs) to achieve high pri-miRNA-binding affinity and specificity (Fig. 1A) (Quick-Cleveland et al. 2014; Nguyen et al. 2015). In the absence of heme, Rhed can still bind RNA (Quick-Cleveland et al. 2014) but Microprocessor displays reduced pri-miRNA cleavage activity biochemically and increased cleavage of pri-miRNAs in the non-productive orientation (Partin et al. 2017; Nguyen et al. 2018). DGCR8 strongly prefers binding Fe(III) heme over Fe(II) heme with over 10^7^-fold difference in affinities, and more importantly only Fe(III) heme can activate DGCR8 for pri-miRNA processing (Barr et al. 2011; Barr et al. 2012). The extraordinary redox specificity of DGCR8 for Fe(III) heme is attributed to a dual-cysteine ligation configuration. DGCR8 coaxially ligates the heme iron using two cysteine side chains, Cys352 from each subunit, in the deprotonated thiolate state, creating an electron-rich environment (Faller et al. 2007; Senturia et al. 2010; Barr et al. 2011). Reduction of the heme iron must induce a conformational change in DGCR8, resulting in loss of activity. We believe that this conformational change involves loss of the cysteine ligands with one or both switched to a different amino acid residue (Barr et al. 2012; Hines et al. 2016), whereas another study suggested protonation of the cysteine thiolates (Girvan et al. 2016).

**Figure 1.**
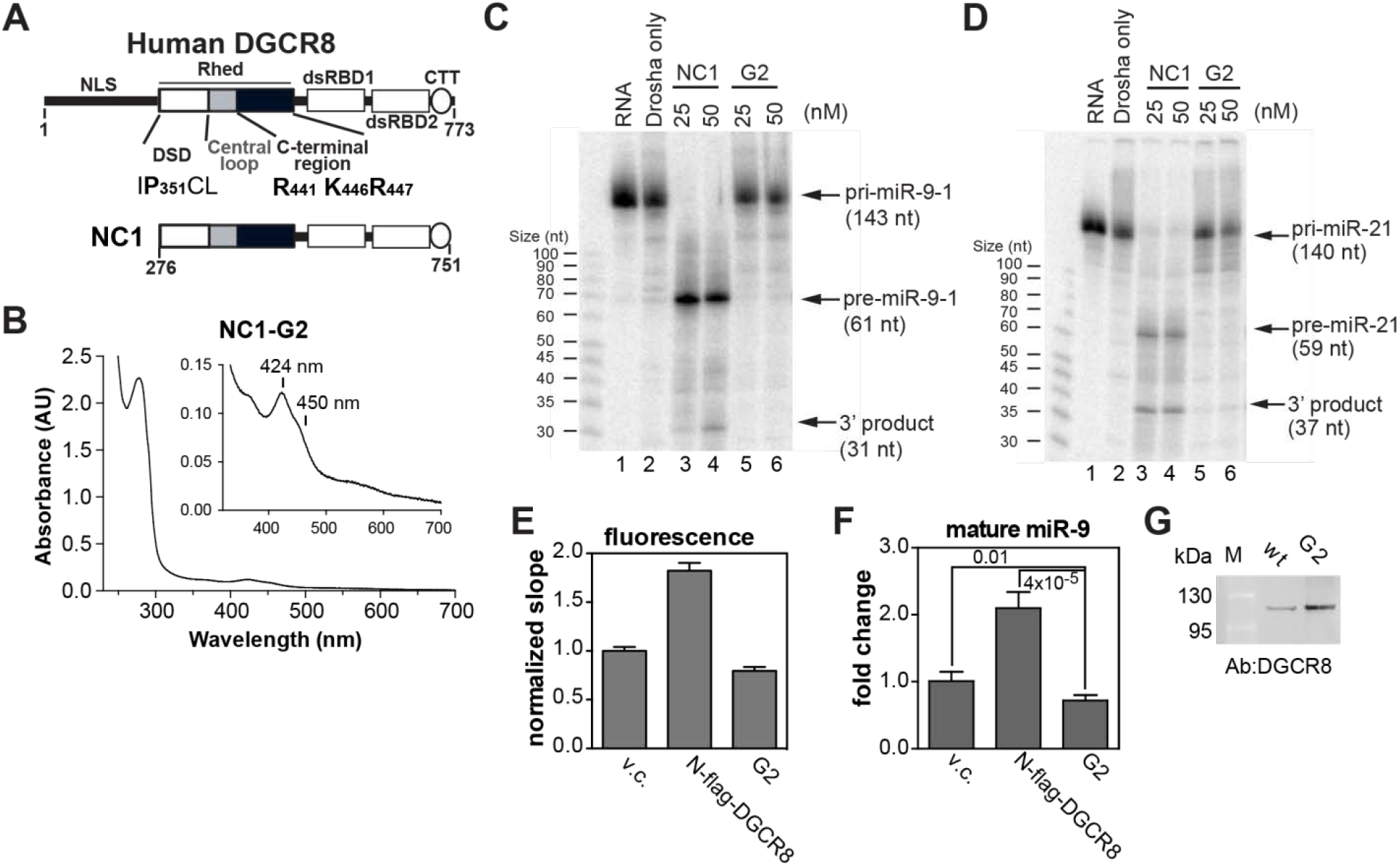
The DGCR8 G2 mutant fails to bind heme and is inactive in pri-miRNA processing. (*A*) Domain structure of human DGCR8, with relevant residues labeled. The NC1 construct is indicated underneath. (*B*) Electronic absorption spectrum of NC1 G2 with an inset showing a zoom-in. No intense 450 nm Soret peak was observed. The protein purity is shown in supplemental Fig. S1. (*C,D*) pri-miRNA processing assays reconstituted using purified Drosha and 25 nM NC1 dimers. (*E*) HeLa cells cultured in complete medium were transfected with pri-miR-9-1 reporter plus either pCMV-tag2a (vector control, v.c.) or N-flag-DGCR8 expression plasmids as indicated. Normalized fluorescence slopes are plotted. Error bars represent 95% confidence interval (CI) from the linear fit. By definition, the fluorescent slopes for which the ranges shown by the 95% CI do not overlap are statistically significant (true for later figures too). (*F*) The abundance of mature miR-9 (mean ± SD, n = 3) normalized by *eYFP* from (*E*) measured using qRT-PCR. *P* values calculated using two-tailed *t*-test are marked on the graph. (*G*) An anti-DGCR8 immunoblot with equal amounts of nuclear extracts loaded.

The specificity of DGCR8 for Fe(III) heme raised the question as to what level of labile Fe(III) heme is necessary to support pri-miRNA processing. We started to address this question by characterizing DGCR8 mutants with various affinities for Fe(III) heme. If the association between DGCR8 and labile Fe(III) heme is under thermodynamic equilibrium in cells, the intracellular labile Fe(III) heme level should be between the dissociation constant (*K*_d_) of the active DGCR8 mutant with the lowest Fe(III) heme affinity and that of the inactive DGCR8 mutant with the highest Fe(III) heme affinity. Cys352 and the surrounding residues, including Ile350, Pro351, and Leu353, constitute the “IPCL” motif that is critical for heme binding (Faller et al. 2007; Weitz et al. 2014). So is Trp329 as part of a WW motif in the dimerization subdomain (DSD) of Rhed (Fig. 1A) (Senturia et al. 2010; Senturia et al. 2012). Mutation of any one of these residues to alanine abolishes DGCR8 activity in our cellular assays (Weitz et al. 2014). Among these mutants, P351A is least defective in heme binding, still capable of binding Fe(III) heme *in vitro* (Barr et al. 2011). The heme-binding affinities of heme proteins may be compared by measuring their dissociation rates (*k*_off_) (Hargrove et al. 1996). The *k*_off_ of P351A-Fe(III) heme complex is around 5 × 10^−4^ s^-1^ at room temperature (Barr et al. 2011). In contrast, the *k*_off_ of WT DGCR8 is too low to be determined as no Fe(III) heme dissociation is observed at all in 4 days (Barr et al. 2011). Based on this evidence, we estimate the *k*_off_ to be ≪3 × 10^−6^ s^-1^, likely ≥1,000 times slower than that of P351A. The large gap between the Fe(III) heme affinities of the active WT DGCR8 and the severely defective P351A mutant (see Results and Discussion for detailed discussion of this mutant’s activity) leaves it uncertain what the labile Fe(III) heme concentration is in cells.

Here we identify DGCR8 mutants that have reduced affinities for Fe(III) heme but maintain their activity. In combination with the severely defective P351A described above, our results reveal a threshold of Fe(III) heme affinity needed for DGCR8 to activate pri-miRNA processing in HeLa cells, reflecting the presence of a labile Fe(III) heme pool. The threshold responds to alterations in cellular heme synthesis and uptake. Based on our understanding of DGCR8 mutants, we offer our interpretation of recently reported miRNA genomic data. We will also discuss mechanistic and functional implications of our findings.

## RESULTS AND DISCUSSION

### The DGCR8 G2 triple mutant does not bind heme and is inactive for pri-miRNA processing

Through systematic mutagenesis of basic residues within the C-terminal region of Rhed, designed for the purpose of another study (Quick-Cleveland et al. 2014), we identified a group mutant G2 (containing R441A, K446A and R447A) that is severely defective in heme binding. The mutations were introduced to NC1, a truncated construct of DGCR8 (a.a. 276-751) that includes all regions biochemically important for pri-miRNA processing activity, namely the Rhed, the dsRBDs and the C-terminal tail (CTT) (Fig. 1A). We overexpressed the mutants in *E. coli*, purified them to homogeneity and examined the presence of heme using electronic absorption spectroscopy. No intense absorption was observed in the 400-500 nm region (Fig. 1B). The minor residual Soret peak was mostly at 424 nm, not the 450 nm characteristic for Fe(III) heme bound to DGCR8 (Fig. 1B inset). These results indicate that one or more of the residues mutated in G2 are important for DGCR8-heme association.

We then measured the activity of the DGCR8 G2 mutant using biochemical and cellular assays. In reconstituted pri-miRNA processing assays, we incubated ^32^P-labeled pri-miRNAs with purified NC1-G2 and His_6_-Drosha^390-1374^ proteins. Denaturing polyacrylamide gel analyses indicated that the pri-miRNA processing activity of G2 was abolished (Fig. 1C,D). To test the cellular activity of G2, we used a fluorescent-based live-cell pri-miRNA processing reporter assay (Weitz et al. 2014). In this assay, a pri-miRNA (pri-miR-9-1) is inserted into the 3′UTR of mCherry-expression cassette so that processing of the pri-miRNA results in degradation of the fusion mRNA and a reduction of mCherry fluorescence. The reporter plasmid simultaneously transcribes *eYFP* and the *mCherry-pri-miRNA* fusion through the use of a bi-directional inducible promoter. The eYFP and mCherry fluorescence signals are measured for individual cells and the slope of eYFP vs mCherry indicates the cellular pri-miRNA processing efficiency. Co-transfection of the reporter with the expression plasmid for full-length DGCR8 (N-flag-DGCR8) increases pri-miRNA processing efficiency, with the fluorescent slope roughly doubled in HeLa cells cultured in complete medium (Fig. 1E). The complete medium contained 10% fetal bovine serum (FBS), the major source of external heme, which brings the heme concentration in the medium to approximately 1 μM. This assay allows us to measure the activity of DGCR8 mutants in cells. Because endogenous DGCR8 is expressed at very low levels in HeLa cells, changes in fluorescence slope mostly reflect the activity of ectopically expressed N-flag-DGCR8 proteins. All inactive DGCR8 mutants characterized in our previous studies give fluorescence slopes similar to that of the vector control (Quick-Cleveland et al. 2014; Weitz et al. 2014). Here we found that expression of N-flag-DGCR8 G2 in HeLa cell cultured in complete medium resulted in a fluorescent slope close to that of the vector control and much lower than WT DGCR8 (Fig. 1E). Quantification of mature miR-9 produced from the reporter supported the conclusion that G2 is inactive in cells (Fig. 1F). Immunoblot analysis showed that G2’s lack of activity was not due to reduced protein expression relative to the WT (Fig. 1G). Altogether, our results suggest that the G2 mutations cause DGCR8 to lose both Fe(III) heme binding and pri-miRNA processing activities.

### G2 single mutants show reduced heme affinity, but remain active in pri-miRNA processing

To determine which residue is responsible for the heme-binding defect of G2, we engineered the individual mutations in the context of NC1 and tested their ability to bind heme when expressed in *E. coli*. As Lys446 and Arg447 are next to each other, we also tested if these two residues function together by characterizing the double mutant K446A/R447A. Similar to G2, the purified NC1 K446A/R447A protein did not contain Fe(III) heme (Fig. S2), suggesting that simultaneous mutation of these residues to a large extent accounts for the heme-binding defect. Somewhat surprisingly, NC1 R441A, K446A, and R447A single mutants were all capable of binding Fe(III) heme (Fig. 2A-C). All three proteins display split Soret peaks around 450 nm and 368 nm, indicating that the double-cysteine ligation is maintained (Barr et al. 2011). One or more of these basic residues may stabilize heme by directly contacting one or both propionate groups, as has been shown in other heme proteins such as myoglobin, human catalase I and hemopexin (Smith et al. 2010).

**Figure 2.**
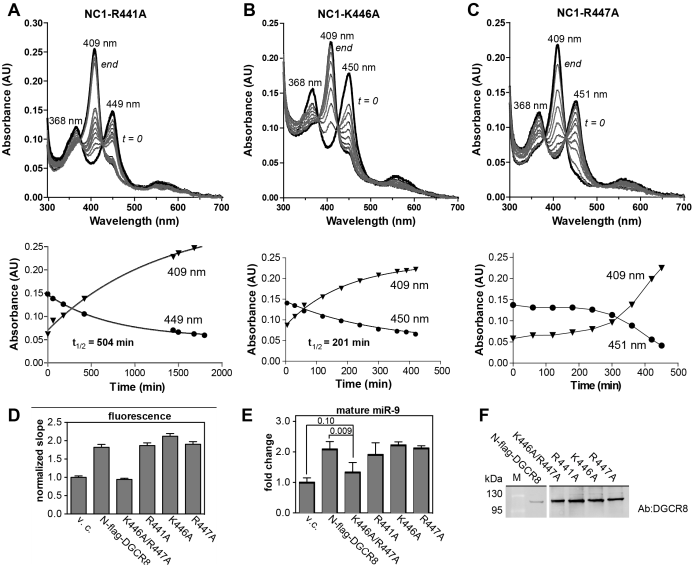
K446A and R447A moderately reduce affinity for Fe(III) heme but remain active in pri-miRNA processing. Fe(III) heme bound NC1 R441A (*A*) K446A (*B*) or R447A (*C*) was incubated with 5 fold excess of apomyoglobin. Top: electronic absorption spectra obtained at different time points. The spectra at time 0 and end are shown by thick black lines. Bottom: *A*_409_ (aquametmyoglobin) and *A*_449/450/451_ (Fe(III) heme-bound NC1) time points were fit to a single exponential (*A, B*) or simply connected by lines (*C*). The *t*_1/2_ values from *A*_449_ or *A*_450_ fits are shown. The purity of these proteins is shown in Fig. S1. (*D*) Cellular pri-miR-9-1 processing assays in complete culture media. Error bars represent 95% CI from the linear fit. (*E*) Abundance of mature miR-9 (mean ± SD, n = 3). *P* values are indicated between pairs of conditions. (*F*) An anti-NC1 immunoblot with equal amounts of nuclear extracts loaded.

We then examined whether the single mutants have reduced heme affinity by measuring their rates of heme dissociation. We incubated the mutants with a five-fold excess of apomyoglobin, which has a high affinity for heme and serves as a heme scavenger. The Fe(III) heme dissociation from DGCR8 is indicated by a reduction in the 450-nm Soret peak and a concurrent appearance of a 409-nm peak indicative of Fe(III) heme transferring to myoglobin. Among the three NC1 mutants, R441A had the slowest Fe(III) heme dissociation, with the half-life (*t*_1/2_) > 8 hr (*k*_off_ = 2.3 × 10^−5^ s^-1^) (Fig. 2A). K446A lost its heme faster with *t*_*1/2*_ ≈ 3.4 hr (*k*_off_ = 5.7 × 10^−5^ s^-1^) (Fig. 2B). The dissociation of heme from NC1-R447A was very slow in the initial 4 hr, but subsequently sped up with most of the Fe(III) heme transferred by the 8th hr (Fig. 2C). The R447A data could not be fit using a single-exponential function, suggesting that the mutant loses heme through a multi-step process. Compared to the previously characterized severely heme-binding-deficient mutations (Faller et al. 2007; Senturia et al. 2010; Weitz et al. 2014), the three G2 single mutations more modestly reduce the affinity for Fe(III) heme.

We subsequently engineered the G2 double and single mutations in the context of N-flag-DGCR8 and measured their cellular activity. In HeLa cells cultured in complete medium, K446A/R447A is inactive in pri-miRNA processing, whereas the three single mutants appear to be as active as WT N-flag-DGCR8, as indicated by both fluorescence slopes and mature miR-9 production (Fig. 2D,E). Immunoblots showed that the mutant proteins were expressed well in HeLa cells (Fig. 2F), ruling out the possibility that the lack of activity for K446A/R447A was caused by reduced expression. Furthermore, similar fluorescence slopes and miR-9 expression levels were obtained when the single mutants were expressed at levels similar to the WT (data not shown). Therefore, the Fe(III) heme affinity of the G2 single mutants must be sufficient to both acquire heme and activate pri-miRNA processing under this experimental condition.

### K446A and P351A reveal a threshold of Fe(III) heme affinity for DGCR8 to be activated

Among the three active G2 single mutants, K446A has the lowest affinity for Fe(III) heme. The *k*_off_ of NC1-K446A is only nine fold slower than that of P351A, the severely defective mutant with the highest affinity for Fe(III) heme known to date. We suggest that the threshold of Fe(III) heme affinity on DGCR8 activity is a reflection of a labile Fe(III) heme pool present in cells. This conclusion depends on the association with heme being primarily responsible for the pri-miRNA processing defects displayed by P351A. We are confident about this assumption, as the mutant’s defects in activity can be rescued by addition of Fe(III) heme biochemically (Fig. S3) and, as will be presented later, by elevation of heme levels in human cells (Fig. 6E,F).

**Figure 3.**
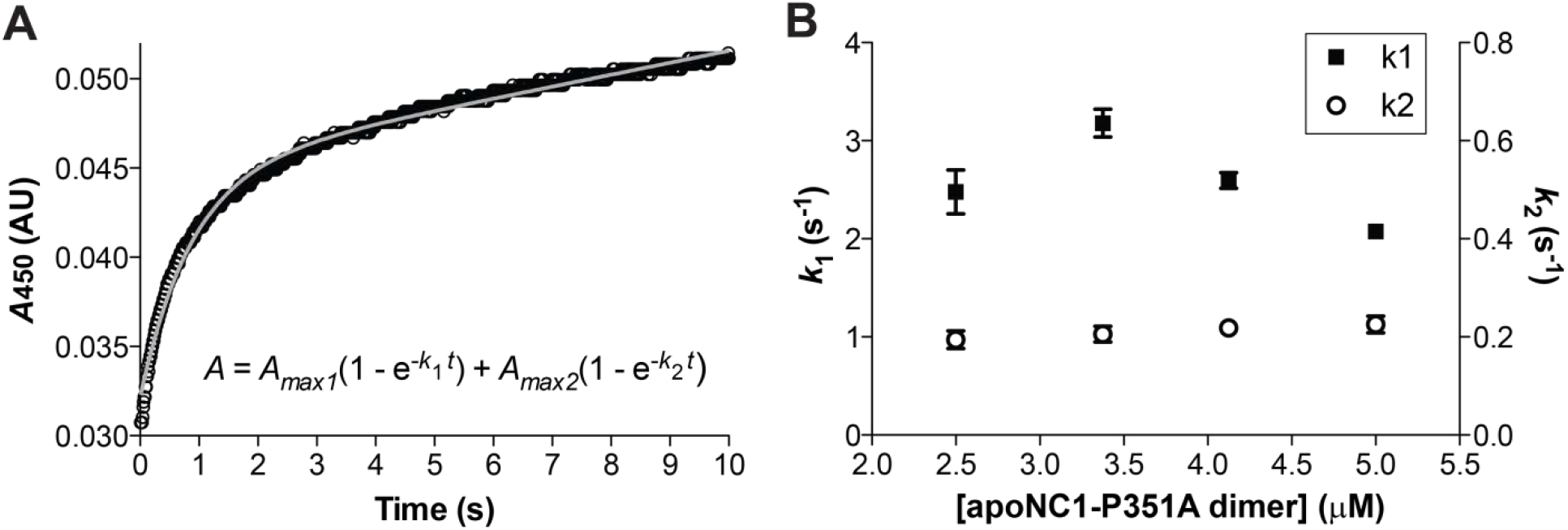
Kinetic analyses of association between apoNC1-P351A dimer and Fe(III) heme. (*A*) A stop-flow *A*_450_ trace of 0.5 μM Fe(III) binding to 4.0 μM apoNC1-P351A dimer. The binding reaction was performed at 25°C in 20 mM Tris/HCl pH 8.0, 400 mM NaCl and 1 mM DTT. The data were fit to the double exponential function shown in the graph where *t* is time, *k*_1_ and *k*_2_ are pseudo-first-order rates, *A*_max1_ and *A*_max2_ are variable during the fit. Individual data points are indicated by circles and the fitted line is in gray. (*B*) The pseudo-first-order rates were measured at four apoNC1-P351A concentrations. Data shown are means ± SD from three or more repeats.

**Figure 4.**
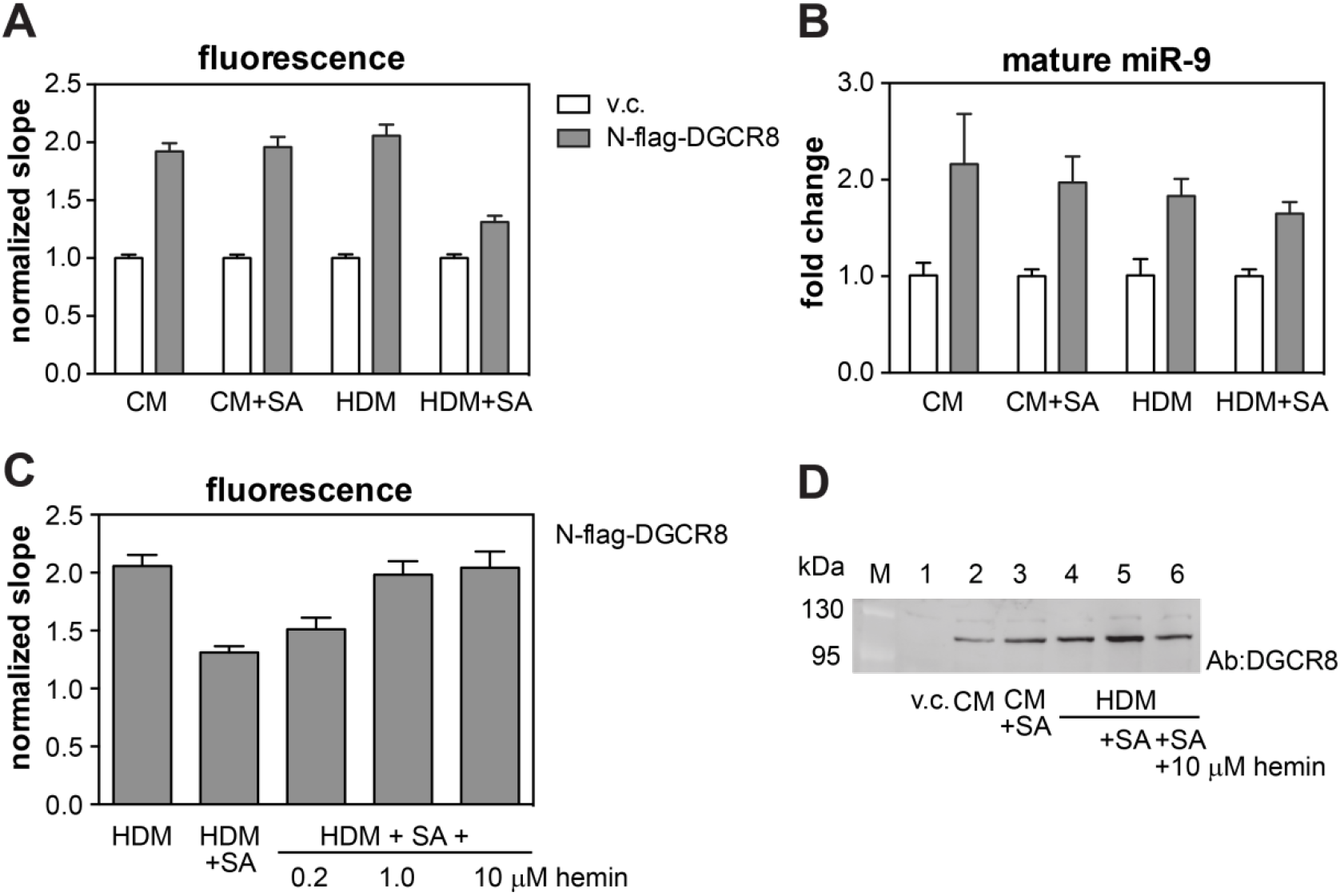
Heme synthesis and uptake influence pri-miRNA processing efficiency of wild-type DGCR8 in HeLa cells. Cellular pri-miR-9-1 processing assays were performed with the N-flag-DGCR8 expression plasmid cotransfected with the reporter. CM: complete media, CM+SA: complete media plus 1 mM succinylacetone, HDM: heme depleted media, HDM+SA: heme depleted media plus 1 mM succinylacetone. (*A*) Normalized fluorescence slopes. Error bars represent 95% CI from the linear fit. (*B*) Abundance of mature miR-9 (mean ± SD, n = 3). (*C*) Cellular pri-miR-9-1 processing assays in which HDM+SA treatment was rescued by adding 0.2, 1, or 10 μM hemin to the media. Error bars represent 95% CI from the linear fit. (*D*) Anti-DGCR8 immunoblots with equal amounts of nuclear extracts loaded.

**Figure 5.**
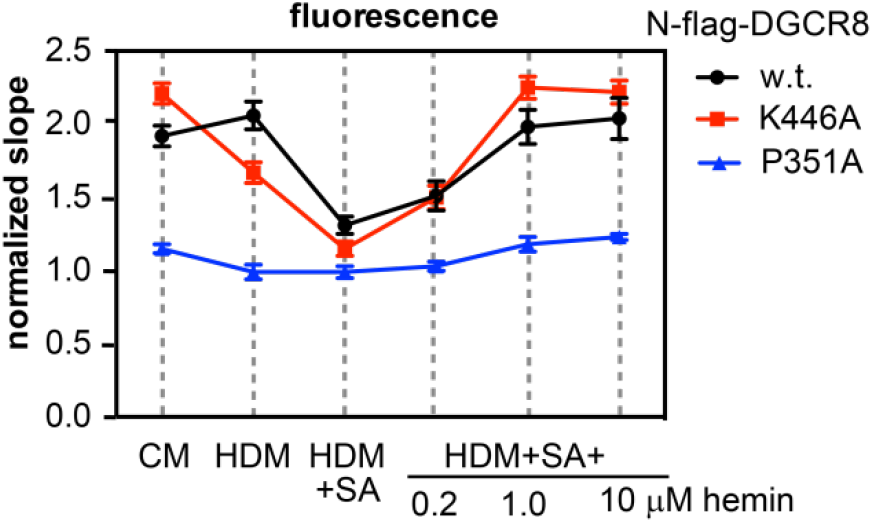
Overall heme level influences the labile Fe(III) heme pool. Cellular pri-miR-9-1 processing assay for N-flag-DGCR8 WT, P351A and K446A under varying heme conditions as described in Fig. 4.

**Figure 6.**
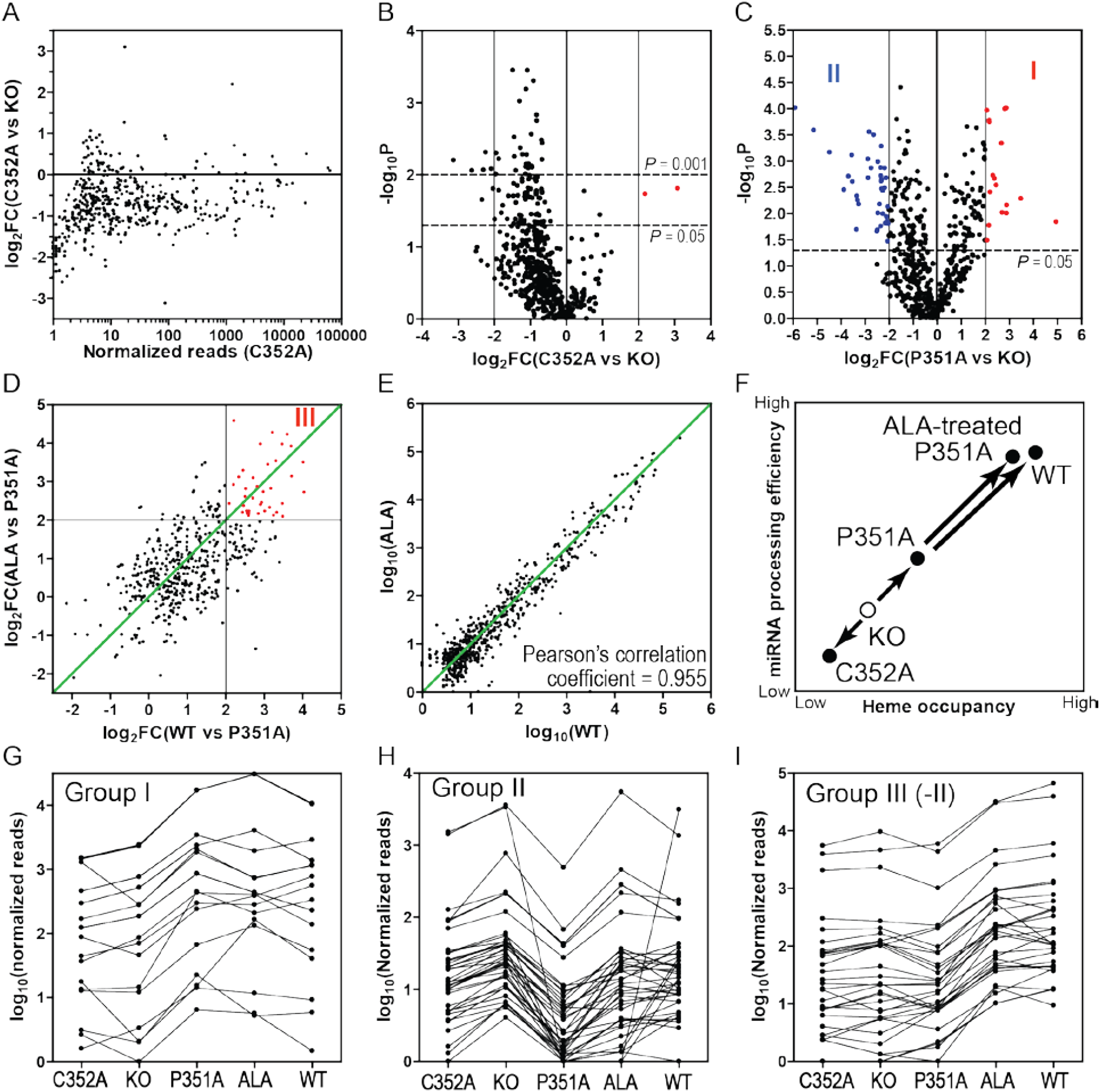
Requirement of heme for miRNA maturation and hierarchy of heme sensitivity among miRNAs. We provide an alternative and deeper interpretation of the next-generation miRNA sequencing data by Kim and colleagues (Nguyen et al. 2018). In this interpretation, we attribute the phenotype of the DGCR8 C352A mutant to its severe defect in binding heme, instead of the combination of defects in both heme binding and dimerization. (A) Distribution of log_2_ fold changes of C352A vs KO over the C352A normalized reads. (B) Volcano plot of the C352A vs KO data sets indicates that most increases of the small number of miRNAs in C352A are not statistically significant. (C) Volcano plot comparing miRNA reads in *DGCR8* KO cells with and without overexpressing the P351A mutant. We identified two sets of miRNAs whose maturation is either enhanced or suppressed by P351A (group I and II, highlighted in red and blue respectively, listed in Table 1). A group II miRNA miR-373 is not displayed on the volcano plot because of its extreme values of log_2_FC at −11.7 and –log_10_P at 4.85. (D) The *y* axis plots log_2_ fold changes of miRNA reads in *DGCR8* KO cells overexpressing the P351A mutant with and without the addition of the heme biosynthesis precursor ALA. The *x* axis plots log_2_ fold changes of miRNA reads in *DGCR8* KO cells overexpressing WT DGCR8 or the P351A mutant. We classify the miRNAs significantly by ALA treatment in P351A-expressing cells and by overexpressing WT DGCR8 as group III (Table 1). (E) Double log_10_ plot of the miRNA reads comparing *DGCR8* KO cells either overexpressing WT DGCR8 cultured in normal medium or overexpressing P351A treated with ALA. The green diagonal line indicates expected position of the miRNAs for which the increases in maturation efficiencies by overexpressing WT DGCR8 and by adding ALA (both relatively to P351A overexpression) are identical. (F) Schematic showing our interpretation, which includes a strong correlation between heme occupancy of DGCR8 and miRNA processing efficiency in cells. KO is defective for reasons not directly related to binding heme and is arbitrarily positioned along the *x* axis. (G,H,I) Comparing the normalized reads of group I, II, and III miRNAs. Some group II miRNAs are also part of group III and are excluded in the plotting for clarity.

### Estimation of Fe(III) heme affinities for DGCR8 variants

We attempted to estimate the affinity of K446A and P351A for Fe(III) heme by measuring their Fe(III) heme kinetic association rate *k*_*on*_ (as *K*_d_ = *k*_off_ / *k*_on_). We previously reported that purified heme-free (apo) NC1-P351A dimer can bind Fe(III) heme to reconstitute a complex with an absorption spectrum very similar to that of the native complex (Barr et al. 2011). In these measurements, we monitored over time the growth of the Soret peak at 447 nm as an indicator of complex formation. Because this binding reaction went to completion quickly, we employed a stopped-flow apparatus. The data were best fit to a two-phase association curve, giving two pseudo-first-order rates *k*_1_ and *k*_2_ (Fig. 3A). To obtain the rate for this second-order reaction, we measured the association rates at four apoNC1-P351A concentrations, all in large excess of the Fe(III) heme concentration. The pseudo-first-order rates should have a linear relationship with the apoNC1-P351A concentration and the slope gives *k*_on_. However, both *k*_1_ and *k*_2_ remained roughly unchanged over the apoNC1-P351A concentration range (Fig. 3B). This is most likely because heme tends to stack with each other in aqueous solutions, forming a variety of oligomers. Unstacking of heme from oligomers is likely to be the rate-limiting step in the kinetic binding reactions. Imidazole and detergents can unstack Fe(III) heme to produce monomers, but they interfere with the DGCR8-Fe(III)-heme interaction. We have not been able to accurately measure the second-order *k*_on_ for DGCR8 and monomeric Fe(III) heme. Nevertheless, our measurements provide a lower *k*_on_ limit of 1 × 10^6^ M^-1^ s^-1^ (maximal k_1_ / [apoNC1-P351A] = 2.5 s^-1^ / 2.5 μM), indicating that association between DGCR8 and Fe(III) heme is very fast.

**Table 1.**
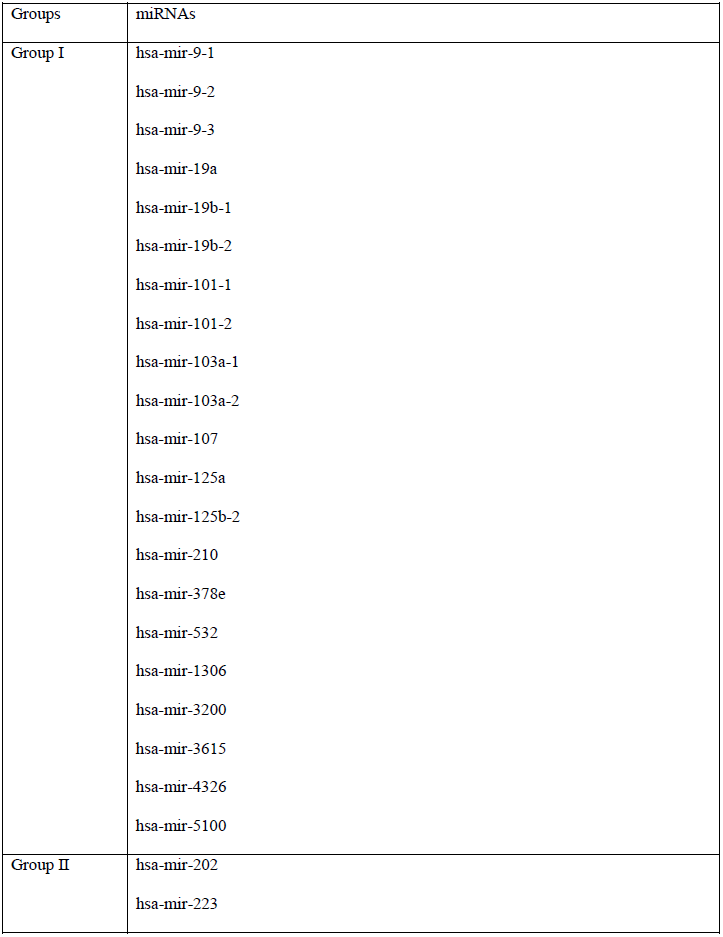
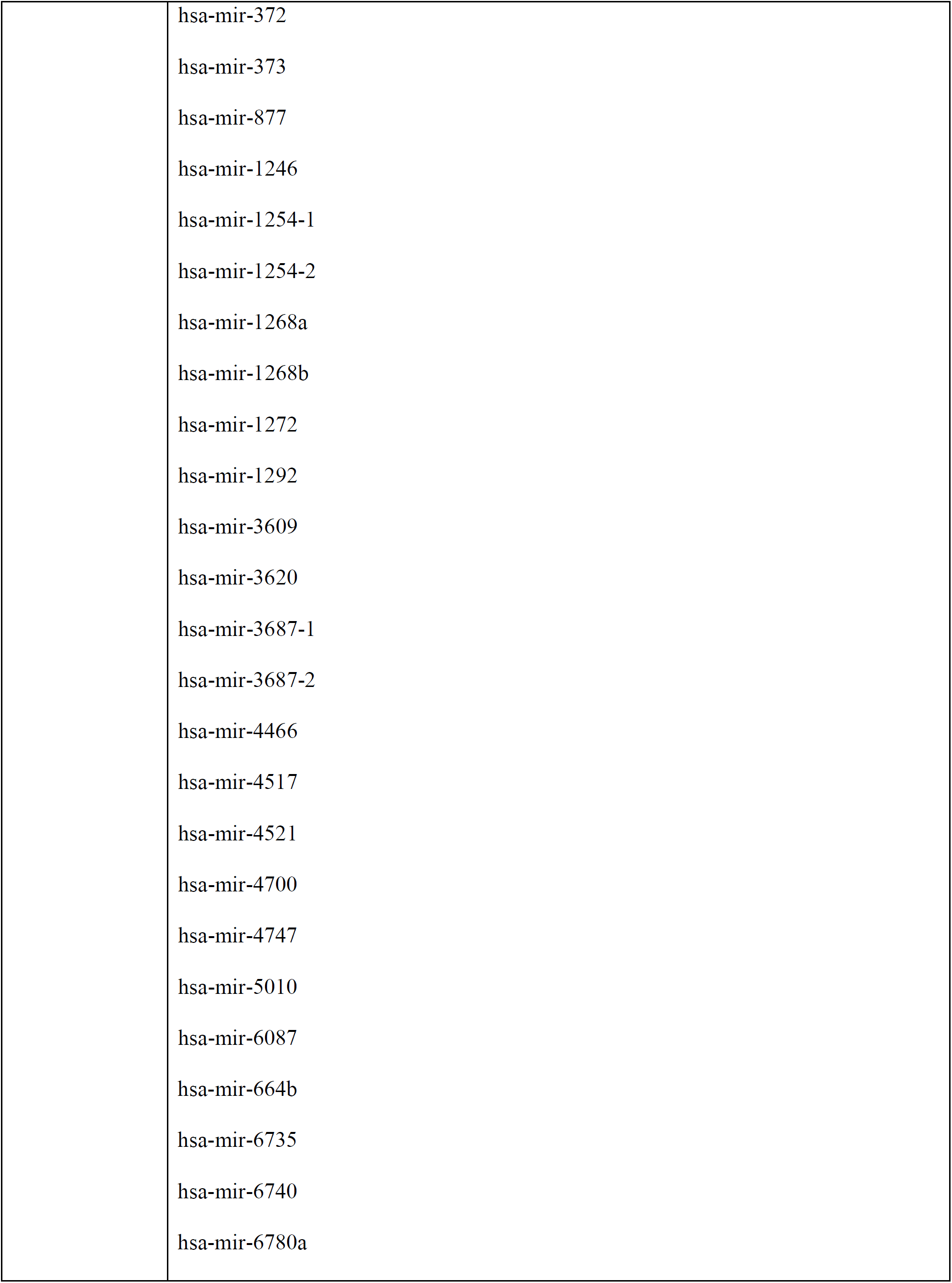
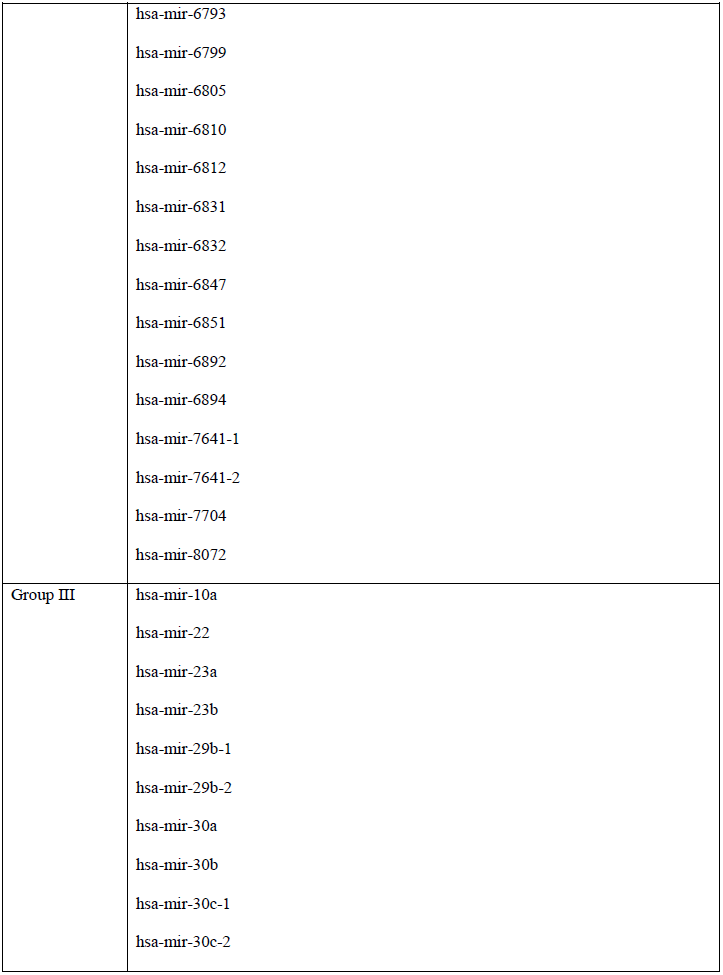
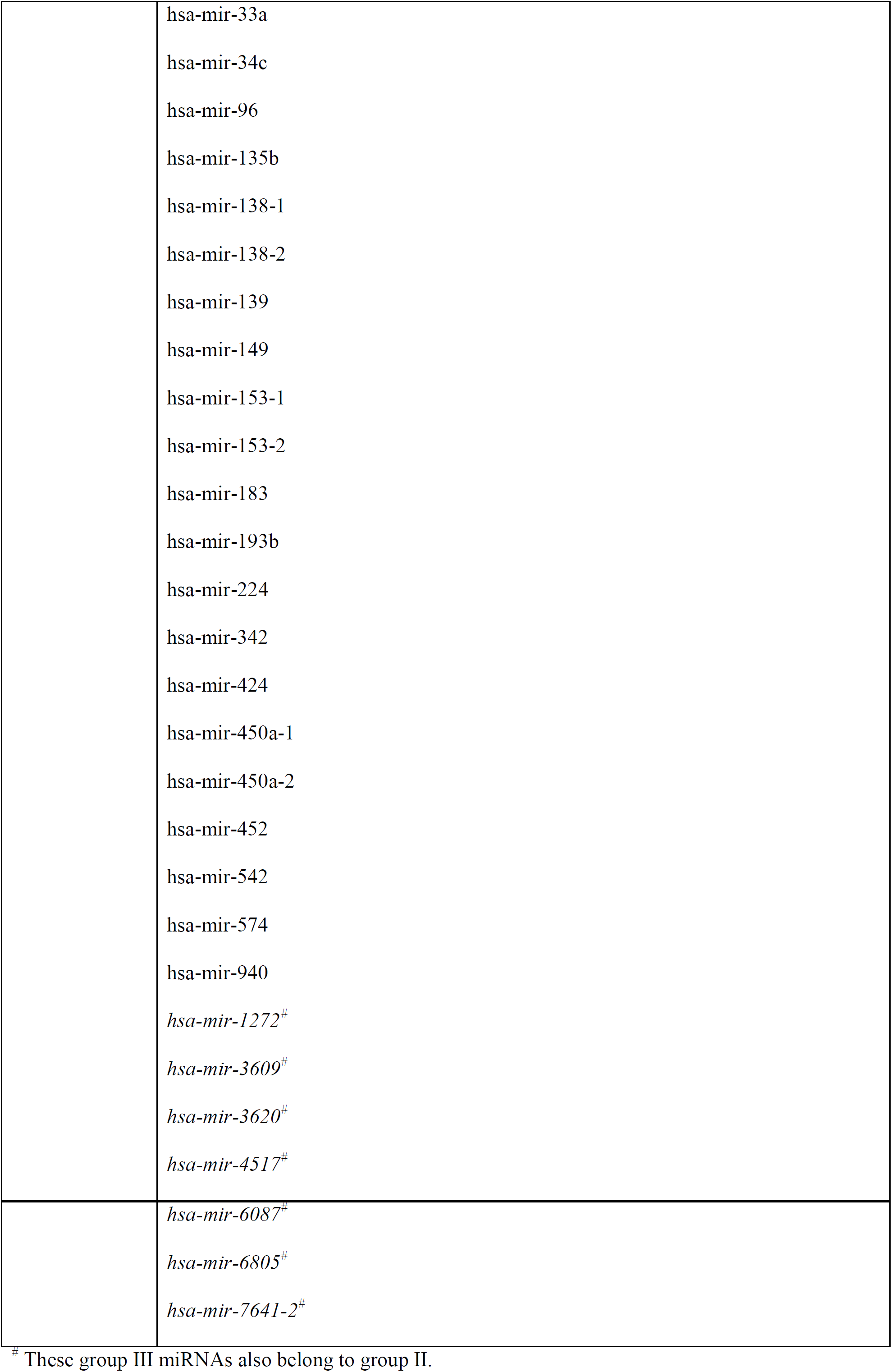
Classification of miRNAs that are differentially influenced by heme-DGCR8 association.

| Groups | miRNAs |
| --- | --- |
| Group I | hsa-mir-9-1<br>hsa-mir-9-2<br>hsa-mir-9-3<br>hsa-mir-19a<br>hsa-mir-19b-1<br>hsa-mir-19b-2<br>hsa-mir-101-1<br>hsa-mir-101-2<br>hsa-mir-103a-1<br>hsa-mir-103a-2<br>hsa-mir-107<br>hsa-mir-125a<br>hsa-mir-125b-2<br>hsa-mir-210<br>hsa-mir-378e<br>hsa-mir-532<br>hsa-mir-1306<br>hsa-mir-3200<br>hsa-mir-3615<br>hsa-mir-4326<br>hsa-mir-5100 |
| Group II | hsa-mir-202<br>hsa-mir-223 |

|  |
| --- |
| hsa-mir-372 |
| hsa-mir-373 |
| hsa-mir-877 |
| hsa-mir-1246 |
| hsa-mir-1254-1 |
| hsa-mir-1254-2 |
| hsa-mir-1268a |
| hsa-mir-1268b |
| hsa-mir-1272 |
| hsa-mir-1292 |
| hsa-mir-3609 |
| hsa-mir-3620 |
| hsa-mir-3687-1 |
| hsa-mir-3687-2 |
| hsa-mir-4466 |
| hsa-mir-4517 |
| hsa-mir-4521 |
| hsa-mir-4700 |
| hsa-mir-4747 |
| hsa-mir-5010 |
| hsa-mir-6087 |
| hsa-mir-664b |
| hsa-mir-6735 |
| hsa-mir-6740 |
| hsa-mir-6780a |
|  | hsa-mir-6793<br>hsa-mir-6799<br>hsa-mir-6805<br>hsa-mir-6810<br>hsa-mir-6812<br>hsa-mir-6831<br>hsa-mir-6832<br>hsa-mir-6847<br>hsa-mir-6851<br>hsa-mir-6892<br>hsa-mir-6894<br>hsa-mir-7641-1<br>hsa-mir-7641-2<br>hsa-mir-7704<br>hsa-mir-8072 |
| Group III | hsa-mir-10a<br>hsa-mir-22<br>hsa-mir-23a<br>hsa-mir-23b<br>hsa-mir-29b-1<br>hsa-mir-29b-2<br>hsa-mir-30a<br>hsa-mir-30b<br>hsa-mir-30c-1<br>hsa-mir-30c-2 |

|  |
| --- |
| hsa-mir-33a |
| hsa-mir-34c |
| hsa-mir-96 |
| hsa-mir-135b |
| hsa-mir-138-1 |
| hsa-mir-138-2 |
| hsa-mir-139 |
| hsa-mir-149 |
| hsa-mir-153-1 |
| hsa-mir-153-2 |
| hsa-mir-183 |
| hsa-mir-193b |
| hsa-mir-224 |
| hsa-mir-342 |
| hsa-mir-424 |
| hsa-mir-450a-1 |
| hsa-mir-450a-2 |
| hsa-mir-452 |
| hsa-mir-542 |
| hsa-mir-574 |
| hsa-mir-940 |
| *hsa-mir-1272<sup>#</sup>* |
| *hsa-mir-3609<sup>#</sup>* |
| *hsa-mir-3620<sup>#</sup>* |
| *hsa-mir-4517<sup>#</sup>* |
|  | *hsa-mir-6087*<sup>#</sup><br>*hsa-mir-6805*<sup>#</sup><br>*hsa-mir-7641-2*<sup>#</sup> |

<sup>#</sup> These group III miRNAs also belong to group II.

Previous characterization of many heme proteins revealed that their *k*_on_’s for heme are very similar and thus the *K*_d_ values are largely determined by the *k*_off_ (Hargrove et al. 1996; Bhakta and Wilks 2006; Nygaard et al. 2006; Liu et al. 2008; Gaudin et al. 2011; Owens et al. 2012). The *k*_on_’s for 35 globins, BSA, and bacterial heme uptake proteins IsdA and Rv0203 are all about 1 × 10^8^ M^-1^ s^-1^, 100-fold higher than the lower *k*_on_ limit we obtained from the stopped-flow experiments. Assuming DGCR8 binds Fe(III) heme with a similar *k*_on_, we estimate that the *K*_d_ of P351A for Fe(III) heme is ∼5 pM, whereas that for K446A is ∼0.6 pM. Therefore, we estimate the threshold of Fe(III) heme affinity to be between 0.6 and 5 pM, as required by DGCR8 to support pri-miRNA processing in HeLa cells cultured in complete medium.

### The DGCR8 activation threshold is strongly influenced by cellular heme synthesis and uptake

We subsequently investigated how heme synthesis and uptake influence the threshold and the labile Fe(III) heme pool. We used four different heme conditions, including (1) the complete medium (CM), (2) complete medium containing 1 mM heme synthesis inhibitor succinylacetone (CM+SA), (3) heme-depleted medium (HDM) in which the heme in serum is partially (∼50%) removed, and (4) heme-depleted medium containing 1 mM SA (HDM+SA). The pri-miRNA processing activity of wild-type N-flag-DGCR8 was not sensitive to partial heme depletion from the medium or inhibition of endogenous heme synthesis alone (Fig. 4A). However, a more severe heme deficiency, caused by combination of the two treatments, rendered the activity of N-flag-DGCR8 substantially reduced (Fig. 4A). Measurements of mature miR-9 produced were generally consistent with the fluorescence slopes, although the larger errors limited further insight (Fig. 4B). The miRNA quantification experiments were likely to be hampered by the coupling of Microprocessor activity with cell growth (Peric et al. 2012).

We were able to rescue the pri-miRNA processing defect under the HDM+SA condition by adding increasing concentrations of hemin to the media (Fig. 4C). A full rescue was achieved at 1 and 10 μM hemin. Even 0.2 μM hemin was able to slightly but significantly rescue the defect. The dose-dependent rescue excluded the possibility that the reduced N-flag-DGCR8 activity is an unintended effect of depleting heme from the serum or the heme biosynthesis inhibitor. Anti-DGCR8 immunoblotting ruled out another possibility that reduced N-flag-DGCR8 expression level is responsible for the pri-miRNA processing deficiency (Fig. 4D). On the contrary, the expression of N-flag-DGCR8 was increased when heme synthesis and/or uptake are inhibited. This observation is consistent with the feedback regulation of *DGCR8* mRNA stability by the Microprocessor activity (Han et al. 2009; Triboulet et al. 2009). Similar effects, but to smaller extents, have been observed in HeLa cells without ectopic expression of N-flag-DGCR8 (Weitz et al. 2014). Altogether, these results suggest that in HeLa cells cultured in complete medium the labile Fe(III) heme pool is not limiting pri-miRNA processing and that heme deficient conditions can deplete the labile Fe(III) heme pool and in turn reduce miRNA maturation efficiency.

In our model, the affinity of K446A for Fe(III) heme is close to the threshold of DGCR8 Fe(III) heme affinity requirement in HeLa cells cultured in complete medium. Therefore, we predict that the activity of this mutant should be more sensitive to heme depletion than WT. Indeed, in the cellular assays for N-flag-DGCR8 K446A, the heme-depleted medium alone caused a large reduction of the fluorescence slope, from 2.21 ± 0.07 (95% CI, same below) in CM to 1.67 ± 0.07 in HDM (Fig. 5). Combination of HDM and SA treatments further deactivated K446A, with the fluorescence slope decreased to 1.15 ± 0.05, also lower than that (1.31 ± 0.06) of WT (Fig. 5). These changes are likely to be direct effects of altering labile Fe(III) heme pool, as addition of hemin to the media rescued the K446A activity in a dose-dependent manner.

In a parallel experiment, we tested the N-flag-DGCR8 P351A mutant. Predictably, P351A remained severely defective under all conditions tested (Fig. 5). We tried to rescue the P351A defect by titrating in 0.2, 1, and 10 μM hemin. Indeed, when 10 μM hemin was added to the medium, the fluorescence slope for P351A increased modestly from 0.99 ± 0.04 (HDM+SA) to 1.23 ± 0.02 and this increase was statistically significant (*p* < 0.001). It is likely that the presence of 10 μM hemin in the medium raised the intracellular labile Fe(III) heme levels so that P351A was able to partially associate with heme. Overall, this series of experiments clearly indicates that the heme synthesis and uptake influence the labile Fe(III) heme pool, which in turn governs DGCR8 activity.

### Genomic evidence supports the requirement of heme for miRNA maturation

We are interested in understanding how DGCR8 heme-binding-deficient mutations and cellular heme conditions affect miRNA maturation at the genomic scale. Recently, Kim and colleagues reported such experiments (Nguyen et al. 2018). They first knocked out *DGCR8* in HCT116, a human colon cancer cell line, by deleting segments of CTT. CTT has been shown to be required for interaction with Drosha (Kwon et al. 2016). The DGCR8 mutants C352A and P351A were then overexpressed and their activities were examined by quantifying the expression of mature miRNAs using next-generation sequencing. They found that in the knockout (KO) cells, the abundance of most canonical miRNAs was greatly reduced whereas their levels relative to each other were maintained. The latter observation suggests that there may be residual miRNA processing activity in the KO cells. Overexpression of C352A caused the abundance of most miRNAs to further decrease, but for some miRNAs the numbers of reads increased relative to those in the KO cells. They attributed the severe phenotype of C352A to its dual defects in dimerization and association with heme.

We analyzed the Kim data set and would like to offer alternative and further interpretations. The first is an alternative interpretation to the C352A phenotype, which provides insight to whether and to what extent heme is required for miRNA maturation. Over 12 years ago, our group reported that recombinant DGCR8 protein expressed in *E. coli* elutes in size-exclusion chromatography as both heme-bound dimer and heme-free monomer and that heme-binding deficient mutations (such as C352A) produce mostly heme-free proteins eluting as monomers (Faller et al. 2007). Based on these observations, we proposed that heme regulates pri-miRNA processing by modulating the oligomerization states of DGCR8. However, our later investigation casted doubt on this model. First, in addition to the intact DGCR8 protein, the “monomeric” species always contain small heterogeneous polypeptides (<10 kDa) that are too small to be resolved by standard SDS 12%-polyacrylamide gels but can by 15% gels (Barr et al. 2011). These small polypeptides are not present in the dimer fractions. Our biochemical characterization and crystal structures show that a short region of ∼55 amino acid residues (the dimerization subdomain of Rhed) is sufficient to mediate dimerization, via an extensive interface (Senturia et al. 2010; Senturia et al. 2012). Therefore, the “monomeric” species may actually be a heterodimer in which one subunit has been cleaved by bacterial proteases to produce a small fragment containing the dimerization subdomain attached to an intact subunit. Second, the “monomeric” DGCR8 cannot bind heme to reconstitute dimers. Third and most importantly, the DGCR8 dimer binds and dissociates with heme reversibly, and the association with Fe(III) heme activates the dimer (Barr et al. 2011; Barr et al. 2012). Based on these observations, we suggest that in human cells in the absence of proteolytic cleavages heme-free DGCR8 proteins, including WT and the mutants, are dimeric.

In our new model, the phenotype of C352A is caused by a near complete loss of association with heme due to its severe heme-binding defect. We compared the miRNA sequencing data of *DGCR8* KO cells with and without overexpressing C352A and found that the small number of miRNAs with increased sequencing reads upon C352A expression are nearly all weakly expressed (Fig. 6A). Most of these increases are small and not statistically significant (Fig. 6B). On the contrary, the expression of most miRNAs are decreased, suggesting that C352A decreases the residual miRNA processing activity in the KO cells and shows dominant negative effects. Therefore, we suggest that C352A is inactive in processing of most if not all canonical pri-miRNAs and thereby heme is required for their maturation.

### Heme modulates the hierarchy and specificity for processing pri-miRNAs

We further analyzed the Kim data by plotting them in various ways. In cells overexpressing P351A, miRNAs have increased, largely unchanged, or decreased expression as compared to the KO (Fig. 6C) (Nguyen et al. 2018). Increasing DGCR8 heme occupancy by either treating the P351A-expressing cells with a heme biosynthesis precursor 5-aminolevulinic acid (ALA) or overexpressing WT DGCR8 causes differential upregulation of most miRNAs (Fig. 6D). The miRNA expression profile of P351A-expressing cells treated with ALA is very similar to that with WT DGCR8 overexpression (Fig. 6E), indicating that the increased heme synthesis and availability rescue the miRNA processing defects of P351A almost entirely. This is in contrast to the slight increase in pri-miRNA processing efficiency of P351A by the addition of hemin observed in our live-cell reporter assays (Fig. 5). This difference may be caused by the differences in the cells (the HCT116 in the Kim study may have higher heme level than the HeLa cells we used), cell culture conditions, or the compounds used (hemin vs ALA, the latter boosts endogenous heme synthesis). Overall, Kim’s study and our data both support the conclusions that P351A reduces the miRNA processing efficiency of DGCR8 primarily by reducing the affinity and occupancy of heme and that labile Fe(III) heme availability in cultured mammalian cells may be close to the ability of P351A to bind Fe(III) heme.

In our interpretation, the phenotypes of both C352A and P351A mutants are caused primarily by their differing degrees of deficiency in binding heme (Fig. 6F). C352A cannot bind heme, whereas without the addition of ALA, overexpression of P351A results in mostly heme-free proteins that are inactive in miRNA processing and exert a dominant negative effect in the KO background. A fraction of P351A should be occupied by heme, allowing maturation of some miRNAs. Therefore, we suggest that Kim’s series of heme experiments mimics conditions under which the Microprocessor complex contains no heme (C352A), partially occupied by heme (P351A), and more substantially (maybe near fully) bound by heme (P351A overexpression with the addition of ALA and overexpression of WT).

Kim and colleagues rightfully concluded that heme differentiates miRNA maturation and their study focused on how heme induces the processing of pri-miRNAs containing a UGU motif at their apical junctions (Nguyen et al. 2018). With new understanding of the mutational effects on DGCR8 activity, we have uncovered previously unknown principles governing how heme affects miRNA maturation. P351A expression mimics a condition in which only a fraction of DGCR8 is occupied by heme. Using a volcano plot (Fig. 6C), we identified two groups of miRNAs, the expression of which is significantly and substantially altered in *DGCR8* KO cells upon P351A expression. Group I contains 21 miRNAs that are enhanced by at least 4 fold (Table 1), indicating that they are preferentially processed when the DGCR8-heme complex concentration is low. For all these miRNAs, the *P* values calculated using *t* test are less than 0.05 and those calculated using Mann-Whitney rank sum test are of the lowest possible values (given only three repeats are available for each condition). ALA treatment or overexpression of WT DGCR8 did not cause large increases of group I miRNAs except miR-1306, which is increased ∼10 fold by ALA treatment (Fig. 6G). Group II contains 42 miRNAs that are suppressed at least 4 fold by P351A expression, with *t*-test *P* values less than 0.05 and lowest possible Mann-Whitney *P* values. In contrast to group I, ALA treatment or overexpression of WT DGCR8 significantly increases the expression of most group II miRNAs (37 out of 42 have *t*-test *P* values less than 0.05) (Fig. 6H). These results suggest that a function of heme association with Microprocessor is to modulate a hierarchy of pri-miRNAs for maturation. When heme is limited, miRNAs like those in group I are still efficiently processed whereas those like in the group II are suppressed. When the heme occupancy is enhanced, the expression of group II is increased whereas that of group I remains unchanged.

Two prominent features distinguish group I and group II miRNAs. First, group I miRNAs have all been characterized and reported by previous studies. Most of them are in the list of 318 “empirical miRNAs” that had been experimentally validated or are conserved, compiled by Lu and colleagues three years ago (Roden et al. 2017). In contrast, most group II miRNAs have not been characterized and only 5 out of 42 are listed in empirical miRNAs. This observation suggests that heme association with Microprocessor prioritizes the maturation of a set of biologically important miRNAs under conditions in which heme is limited. Second, primary transcripts of group I miRNAs tend to display structural features known to be favorable for processing, whereas those of group II miRNAs often do not. Among the 21 group I miRNAs, 48% fold to hairpins with favorable stem lengths between 33-39 nt and 48% with favorable apical loop size between 8 and 16 nt. Among the 42 group II miRNAs, only 24% have favorable stem lengths and 24% with optimal apical loop sizes. Two (9.5% of 21) group I pri-miRNAs contain UGU motif at the apical junction, whereas the group II has none. These structural and sequence differences explain why the processing of group I pri-miRNAs is prioritized.

It has been suggested that some miRNAs registered in miRBase are not authentic miRNAs (Guo et al. 2015). It is possible that some group II miRNAs are such cases. In this sense, an appropriate level of DGCR8-heme complexes in cells should maximize the specificity of the processing machinery, as biologically important, bona fide miRNAs are efficiently processed, and cleavage of questionable pri-miRNAs or pri-miRNA-like hairpins is suppressed. Together these observations suggest that a function of heme in miRNA maturation is to modulate the specificity of the processing machinery.

### Changes in cellular heme level and DGCR8 heme occupancy alter the expression of a subset of miRNAs

Our previous works (Weitz et al. 2014; Barr et al. 2015) and the data presented here show that changes of cellular heme level modulate pri-miRNA processing efficiency in the contexts of both endogenous WT DGCR8 expression and overexpression of DGCR8 (WT and mutants). Although the physiological conditions in which heme regulates miRNA maturation and how pathological dysregulation alters miRNA processing remain to be determined, the Kim data set provides insight into relevant changes in miRNA expression pattern. This is useful as the principles and underlying molecular mechanisms are likely to be conserved. The fact that treating P351A-expressing KO cells with ALA and expression of WT DGCR8 achieve similar miRNA expression profiles suggests that heme occupancy of DGCR8 is key to determine the activity and specificity of the miRNA processing machinery. The heme occupancy may be elevated by increasing the cellular heme levels or by increasing the affinity of DGCR8 for heme.

From the Kim data set, we identified a set of miRNAs the expression of which changes upon alteration of intracellular heme level or DGCR8’s heme affinity. From the volcano plot (Fig. 6D), we identified 38 miRNAs that are >4-fold up-regulated both by ALA in P351A-expressing cells and by WT overexpression relative to P351A. We call this set of miRNAs group III (Table 1). Importantly, no group III miRNAs increase by more than 4 fold when P351A is expressed compared to KO (Fig. 6I). Only five (mir-224, mir-450a-1, mir-450a-2, mir-452 and mir-542) increase moderately (by 2-4 fold). Therefore, we consider group III miRNAs to be selectively up-regulated when heme occupancy of Microprocessor goes from partial to high. Not surprisingly, group III contains 7 miRNAs that are also in group II, as most group II miRNAs are up-regulated by ALA or by overexpressing WT instead of P351A (Fig. 6H). The remaining miRNAs, we label as group III-II, are mostly well characterized miRNAs, including miR-10a, miR-22, miR-23a,b, miR-29b, miR-30a,b,c, miR-96, miR-183, and miR-193b. All of them have well documented roles in cancer. These observations imply that heme-mediated miRNA expression changes may have a role in carcinogenesis. Furthermore, consistent with their intermediate status in the processing hierarchy, group III-II pri-miRNAs display stem lengths (42% in the favorable 33-39 range) and apical loop sizes (42% in the range of 8 and 16) better than group II (24% for both features) but worse than group I (48% for both). We note that 5 out of 31 (16%) Group III-II pri-miRNAs contain UGU motif at the apical junction, significantly higher (*P* = 0.02) than the frequency of 5.3% among all human pri-miRNAs.

Our results and analyses shed new light to the debate whether heme bound to DGCR8 serves a house-keeping or regulatory function. The large gap between the activation threshold when cells are cultured in complete medium and the dissociation constant of WT DGCR8 for Fe(III) heme favors the house keeping idea, e.g. DGCR8 is loaded with Fe(III) heme all the time to keep the miRNA processing pathway on. This idea is further supported by the fact that it is difficult to biochemically reduce the Fe^3+^ in the heme bound to DGCR8. Long incubation (30 min) with sodium dithionite is required for a full reduction (Barr et al. 2012; Hines et al. 2016). Such reduction is unlikely to occur under physiological conditions. On the other hand, it is premature to rule out the regulatory function, especially as we know very little about labile Fe(III) heme in cells. Heme deficient conditions can clearly reduce miRNA maturation, which may occur under certain biological contexts and certainly in diseases. The realization of differential sensitivity of miRNAs to heme and the classification of miRNAs into corresponding groups provide a basis for further investigation. Operationally, the house-keep and regulatory functions are not mutually exclusive. For example, if labile Fe(III) heme level only changes moderately under normal physiological conditions, heme would be house-keeping for group I miRNAs but might be regulatory for group III miRNAs.

### A potential labile Fe(III) heme pool

Our study suggests the presence of a labile Fe(III) heme pool (Fig. 7). The concentration of the “general” labile heme pool estimated using heme sensors varies from 6.5 nM (Song et al. 2015), 20-40 nM (cytoplasm) and <2.5 (mitochondrion and nucleus) (Hanna et al. 2016), and 433 nM (Yuan et al. 2016). All these values are much higher than the pM Fe(III) heme affinity threshold for activating miRNA maturation we observed. Therefore, the labile Fe(III) heme pool must be distinct from the general labile heme pool, which is thought to be dominated by Fe(II) heme as the cellular compartments are largely reducing (with the exception of endoplasmic reticulum).

**Figure 7.**
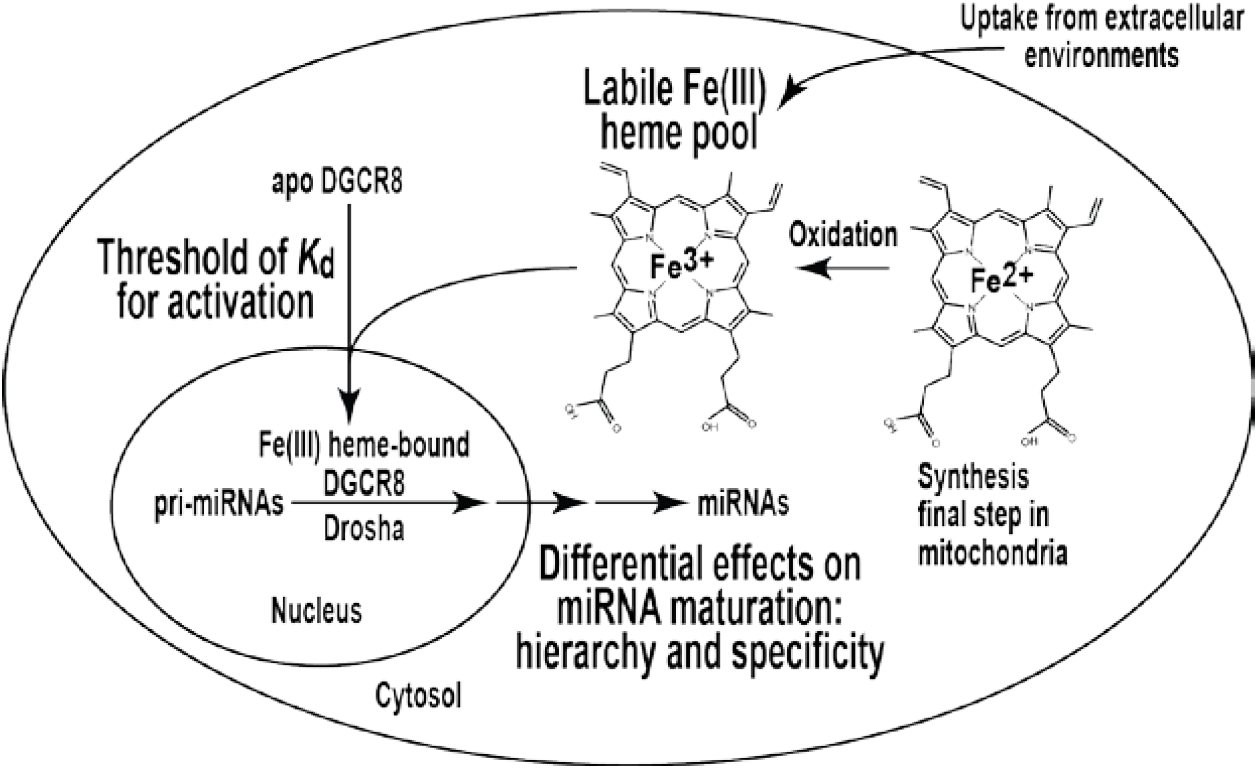
Schematic of Fe(III) heme’s role in miRNA maturation. See main text for detailed explanation.

The high threshold of DGCR8 Fe(III) heme affinity required for activity (0.6-5 pM in terms of *K*_d_ values) intrigued us to think deeper into the status of labile Fe(III) heme in cells. Here we consider two scenarios, labile Fe(III) heme existing without and with a transporter/chaperone protein. If labile Fe(III) heme is free from transporters and chaperones, under thermodynamic equilibrium, the threshold would be simply determined by the concentration of labile Fe(III) heme in cells. Given that the volume of a HeLa cell is typically ∼2,000 μm^3^, this concentration means that only one or a few labile Fe(III) heme molecules (concentration x volume x Avogadro constant = 0.7–6 molecules per cell) exist in each cell cultured in complete medium. This number would be even lower in the smaller volume of the nuclear compartment. The minute number of “free” Fe(III) heme molecules leaves little room for further reduction under various biological and disease conditions, as mimicked by heme depletion and inhibition in our investigation.

It seems more likely that labile Fe(III) heme is carried by a Fe(III) heme-specific transporter or chaperone. In this scenario, the threshold results from a requirement of DGCR8 to receive Fe(III) heme from the transporter/chaperone and there can be many labile Fe(III) molecules per cell, constituting a pool large enough for reliable control of relevant biological processes. Under heme deficient conditions, fewer transporter molecules are loaded with Fe(III) heme and thereby fewer DGCR8 proteins receive heme and are activated for pri-miRNA processing.

### The redox antenna hypothesis

Heme is synthesized exclusively in the Fe(II) form (Dailey 2002) and has to be oxidized to the Fe(III) state before activating miRNA maturation (Fig. 7). The heme iron oxidation may occur during trafficking from mitochondria, the site of synthesis, to its destination, which could be the cytoplasm where the DGCR8 protein is translated or the nucleus where pri-miRNA processing occurs. The conversion of Fe(II) heme to the Fe(III) forms should be influenced by cellular redox environments, including the various redox buffering systems, cellular metabolism, and cell growth states. Redox regulation of labile heme may also be linked to heme degradation, and import and export across cellular membranes (Severance and Hamza 2009; Khan and Quigley 2011). Redox properties of heme are critical to its function as a prosthetic group of proteins. Our study suggests that redox sensitivity of heme is important for its signaling functions. In this sense, labile heme may serve as a “redox antenna”.

Among well-characterized cysteine-ligated heme proteins, DGCR8 is unique in that it simultaneously uses two cysteine side chains as ligands. Such configuration allows its extraordinary preference for Fe(III) heme for binding and activation, which enables us to use live-cell pri-miRNA processing activity as readout to confidently detect labile Fe(III) heme. However, the function of labile Fe(III) heme is unlikely to be limited to miRNA maturation. There is a class of heme proteins that has been proposed to be heme sensors (Shimizu 2012; Girvan and Munro 2013). They use a single cysteine thiolate at a given time for axial ligation, but still generally prefer Fe(III) heme over Fe(II) heme. These sensors include the Rev-erbα and Rev-erbβ nuclear receptors (Raghuram et al. 2007), the Bach1 transcription factor (Ogawa et al. 2001), and the yeast Gis1 dual-function transcription factor and histone demethylase (Lal et al. 2018). They are involved in diverse functions such as circadian rhythm, transcription regulation of metabolic pathways, heme catabolism, ion channel activities and protein degradation. We are only beginning to understand the roles of labile Fe(III) heme in biology and relevant diseases.

## MATERIALS AND METHODS

### Plasmids

Mutagenesis was performed using 4-primer PCR. For bacterial expression of NC1 R341A, K446A, R447A and K446A/R447A, DGCR8-coding sequences were inserted between NdeI and EcoRI sites in the pET-24a+ vector. For mammalian expression of N-flag-DGCR8 mutants R341A, K446A, R447A and K446A/R447A, DGCR8-coding sequences were inserted between BamHI and EcoRI sites in the pCMV-tag2a vector, resulting in a N-terminal flag-tagged DGCR8 protein. The coding sequences in all plasmids were confirmed using sequencing.

### Expression, purification and characterization of DGCR8 (NC1) proteins

Expression, purification and reconstituted pri-miRNA processing assays were done as described (Barr and Guo 2014). Human NC1 mutants were expressed in the presence of 1 mM δ-ALA. Purification was done with cation exchange followed by size exclusion chromatography (SEC). The purified dimeric proteins were stored at 4°C in SEC buffer containing 20 mM Tris pH 8.0, 400 mM NaCl and 1 mM dithiothreitol and were used in characterization within two days. The electronic absorption spectra were recorded using a CARY 300 spectrophotometer (Agilent) with the bandwidth set to 0.5 nm. The reconstituted pri-miRNA processing reactions contained pri-miRNAs uniformly labeled using ^32^P, recombinant Drosha, and purified DGCR8. The reactions were incubated at 37°C for 45 min and were analyzed using 15% polyacrylamide denaturing gels containing 7 M urea and autoradiography.

### Live-cell pri-miRNA processing assays

The assay has been described previously (Weitz et al. 2014). Succinylacetone (MilliporeSigma) and hemin (MP Biomedicals) stock solutions were prepared in water and DMSO, respectively. Cells were grown either in complete medium (DMEM with 10% FBS) or heme-depleted medium and then split into glass-bottom plates for imaging. The heme-depleted medium was the same as the complete medium except that the FBS was treated with 0.1 M ascorbic acid for 2 hr (Ye and Zhang 2004; Weitz et al. 2014). The ascorbic acid was removed via two rounds of dialysis at room temperature against 1 L of phosphate buffered saline (137 mM NaCl, 2.7 mM KCl, 10 mM Na_2_HPO4, and 1.8 mM KH_2_PO4) for 1 hr each round. The heme contents of FBS with and without heme depletion were measured using pyridine hemochromagen assays as described (Barr and Guo 2015). Cells were treated with 1 mM SA either alone or with hemin from 2 hr prior to transfection until imaging. Cells were cotransfected with the pri-miR-9-1 reporter and the indicated N-flag-DGCR8 construct. In the experiments shown in Fig. 2D-2F, we used twice the amount of N-flag-DGCR8 mutant plasmid as the WT in the cotransfections. Immediately after transfection, doxycycline was added to a final concentration of 1 μM to induce reporter expression. Cells were imaged 20-24 hr post-induction. The fluorescent intensities of individual cells in the images were integrated using a MATLAB script as described (Quick-Cleveland et al. 2014; Weitz et al. 2014). The mCherry and eYFP intensities were plotted and fit to a linear regression using PRISM (GraphPad, version 4), in which the statistical analyses were also performed. The accompanying qRT-PCR and immunoblotting were performed as described (Quick-Cleveland et al. 2014; Weitz et al. 2014). In our experience, the eYFP vs mCherry fluorescence slope is not very sensitive to variation in N-flag-DGCR8 overexpression level.

### Heme off-rate measurements

The measurements were performed as previously described (Barr et al. 2011). Horse skeletal muscle apomyoglobin was dissolved in SEC buffer and the concentration was measured by absorption spectrometry (ε_280nm_ = 13,940 M^-1^ cm^-1^). Five-fold molar excess of apomyoglobin was added to NC1 mutants and transfer of heme was measured by conversion of NC1 Soret peak at ∼450 nm to that of myoglobin at 409 nm. Electronic absorption spectra were recorded at room temperature on a Cary 300 spectrophotometer with bandwidth set to 1 nm. Prism software (GraphPad, version 4) was used to fit and plot the absorbance over time. The data were fit with a one-phase exponential decay function to calculate the *k*_off_.

### Analysis of small RNA sequencing data of DGCR8 KO and rescue experiments

The previous reported data were obtained from the Gene Expression Omnibus (GEO: http://www.ncbi.nlm.nih.gov/geo) under the accession number GSE111431 (Nguyen et al. 2018). In the downloaded data, the miRNA reads have already been normalized. All data analyses were performed using Microsoft Excel (version 2010) except the Mann-Whitney tests, which were done in R. In each analysis, we excluded miRNAs with maximal read counts less than 5. To avoid losing miRNAs in fold change and log calculations, a pseudo-count of 1 was added to all read counts. The *P* values shown in the volcano plots were calculated using two-tailed *t* test assuming two-sample equal variance (homoscedastic). The *t* test was performed using log_2_(reads) of the three repeats under each condition as inputs. All plots were generated using PRISM (version 4, GraphPad).

## SUPPLEMENTAL MATERIAL

Supplemental material is available for this article.

## ACKNOWLEDGMENTS

We thank Jason Ernst for advice on genomic data analyses. This work was supported by NIH grant GM080563 and by a seed grant from the Spitzer Foundation to F.G., USPHS National Research Service Award 5T32GM008496 and UCLA Dissertation Year Fellowship to S.H.W., and Ruth L. Kirschstein National Research Service Award GM007185 to J.Q.

**The authors declare no conflict of interest.**

