## Supplemental Figures for "Fe(III) heme sets an activation threshold for processing distinct groups of pri-miRNAs in mammalian cells"

*From the [1] Interdepartmental Program in Molecular, Cellular, and Integrative Physiology, [2] Department of Biological Chemistry, David Geffen School of Medicine, [3] Bioinformatics Interdepartmental Program, [4] Department of Integrative Biology and Physiology, [5] Department of Chemistry and Biochemistry, [6] Department of Physiology, David Geffen School of Medicine, [7] Molecular Biology Institute, University of California, Los Angeles, California, 90095*

*[8] Current affiliation: Department of Molecular Cell & Developmental Biology, University of California Santa Cruz, Santa Cruz, CA 95064*

*[9] Current affiliation: Dominican University of California, San Rafael, CA 94901*

### Supplemental Figures

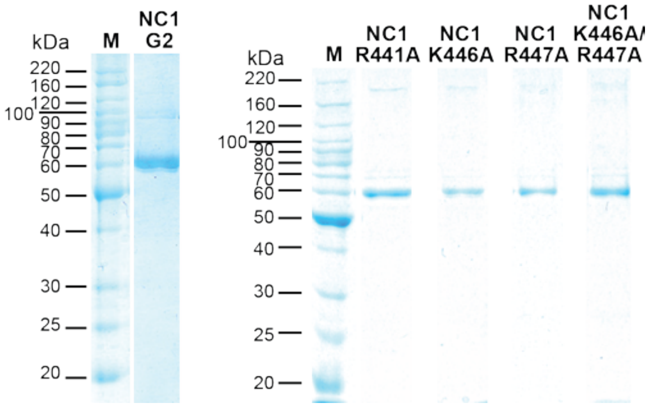


**Figure S1. SDS gel analyses of purified NC1 mutant proteins.**
The proteins were analyzed using 12% SDS gels stained with Coomassie.

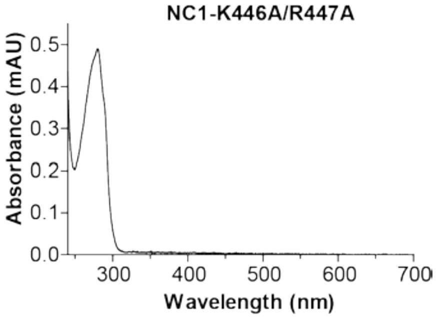


**Figure S2. Electron absorption spectrum of purified NC1 K446A/R447A.**
This protein was expressed in *E. coli* in the presence of 1 mM σ-aminolevulinic acid, and was purified using cation exchange and size exclusion chromatography. The purity is demonstrated in Fig. S1. The absorption spectrum was recorded at room temperature in 20 mM Tris pH 8.0, 400 mM NaCl and 1 mM DTT.

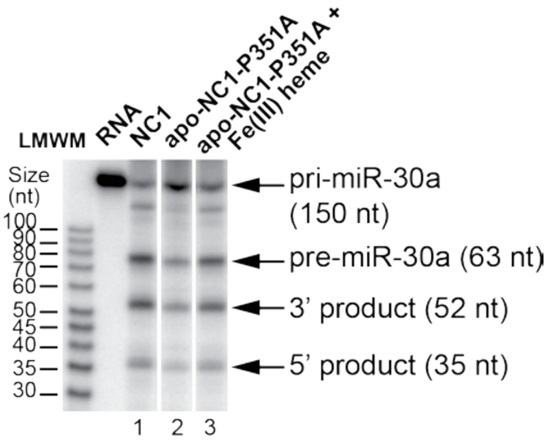


**Figure S3. The P351A mutation does not hamper the Fe(III) heme-mediated activation of pri-miRNA processing.**

In vitro pri-miR-30a processing assays using a trace amount of uniformly $^{32}$P-labeled RNA, recombinant full-length Drosha and various forms of DGCR8 (at 25 nM of dimer), including native Fe(III) heme-bound NC1 (lane 1), apoNC1-P351A (lane 2), and apoNC1-P351A incubated with equimolar of Fe(III) heme (lane 3). The reactions were incubated at 37˚C for 45 min, and were analyzed using a 7M-urea 15% polyacrylamide gel. All lanes shown were from the same gel.
